# The principle of inverse effectiveness in audiovisual speech perception

**DOI:** 10.1101/585182

**Authors:** Luuk P.H. van de Rijt, Anja Roye, Emmanuel A.M. Mylanus, A. John van Opstal, Marc M. van Wanrooij

## Abstract

We assessed how synchronous speech listening and lip reading affects speech recognition in acoustic noise. In simple audiovisual perceptual tasks, inverse effectiveness is often observed, which holds that the weaker the unimodal stimuli, or the poorer their signal-to-noise ratio, the stronger the audiovisual benefit. So far, however, inverse effectiveness has not been demonstrated for complex audiovisual speech stimuli. Here we assess whether this multisensory integration effect can also be observed for the recognizability of spoken words.

To that end, we presented audiovisual sentences to 18 native-Dutch normal-hearing participants, who had to identify the spoken words from a finite list. Speech-recognition performance was determined for auditory-only, visual-only (lipreading) and auditory-visual conditions. To modulate acoustic task difficulty, we systematically varied the auditory signal-to-noise ratio. In line with a commonly-observed multisensory enhancement on speech recognition, audiovisual words were more easily recognized than auditory-only words (recognition thresholds of −15 dB and −12 dB, respectively).

We here show that the difficulty of recognizing a particular word, either acoustically or visually, determines the occurrence of inverse effectiveness in audiovisual word integration. Thus, words that are better heard or recognized through lipreading, benefit less from bimodal presentation.

Audiovisual performance at the lowest acoustic signal-to-noise ratios (45%) fell below the visual recognition rates (60%), reflecting an actual deterioration of lipreading in the presence of excessive acoustic noise. This suggests that the brain may adopt a strategy in which attention has to be divided between listening and lip reading.

## Introduction

Speech is a complex, dynamic multisensory stimulus, characterized by both an auditory and a visual information stream. Congruent information of the sensory modalities (i.e. spatial and temporal coincidence of the sensory streams, and their meanings) is integrated in the brain [1,2] to form a coherent, often enhanced, percept of the common underlying source [3]. Indeed, additional synchronous visual information (i.e. speech-reading/lipreading) has a positive impact on speech perception, and audiovisual speech recognition in acoustic noise is substantially better than for auditory speech alone [4–15].

Audiovisual integration in general, has been the topic of a variety of behavioral and electrophysiological studies, involving rapid eye-orienting to simple peripheral stimuli [16,17], spatial and temporal discrimination of audiovisual objects [18–20], and the integrative responses of single neurons in cats and monkeys [21–23]. Three main principles have been shown to govern the mechanisms of multisensory integration: i. spatial alignment of the different sources, ii. temporal (near-)synchrony, and iii. inverse effectiveness. The latter holds that multisensory enhancement strongly increases for poorly perceptible unisensory signals, for example in the presence of acoustic background noise or visual distracters [3]. Although these principles have mostly been demonstrated at the neurophysiological level of anesthetized experimental animals (for review, see [3]), several studies on audiovisual saccadic eye movements in humans or on manual reaction times in macaques and humans [24], have revealed systematic modulations of the effects of audiovisual congruency and inverse effectiveness that corroborate the neurophysiological data [16,25,26].

In this study, we focus on whether the phenomenon of inverse effectiveness can also be applied to speech perception. This is not a trivial extension of the classical audiovisual integration studies, as the underlying speech-related sensory signals are complex and dynamic signals, requiring advanced (top-down) neural processing within the auditory and visual systems. One way of studying the presence of inverse effectiveness in the perception of audiovisual speech stimuli is by adding background noise [11,15,27], which effectively changes the saliency of the auditory stimulus. By doing so, earlier studies have suggested an absence of inverse effectiveness, as at low unimodal performance scores, the audiovisual enhancement decreases. The principle of inverse effectiveness has also been studied by quantifying the differences in unimodal word-recognition performance scores across (groups of) subjects [7,15,28,29], however, outcomes were not consistent. To our knowledge, the effect of the visual or auditory recognizability of words (irrespective of background noise) on the presence or absence of inverse effectiveness has not been studied. For example, if certain words would be simply more salient than other words, these might be better heard or visually recognized over a large range of noise levels. If the principle of inverse effectiveness would hold at the word level, the highly-salient words should benefit less from bimodal presentation than the less salient words. To study this possibility, we determined how well words can be recognized by listening and/or lipreading under noisy listening conditions in normal-hearing subjects.

## Results

### Overview

Eighteen normal-hearing subjects had to identify 50 words occurring in 155 unique five-word sentences, by selecting the words they recognized (ten-alternative forced choice) on a screen. The speech material was based on the Dutch version of the speech-in-noise matrix test developed by Houben and colleagues [30] (see Methods on the construction of the speech material). The words were presented in acoustically-only (A-only), visual-only (V-only) or bimodal (AV) blocks. An acoustic background noise was played in the A-only and AV conditions at five signal-to-noise ratios.

### Lip reading

We will first describe the lipreading abilities (V-only). These were quantified for every subject (n=18) and every word (n=50) as the number of correct responses, z, divided by the number of presentations, N, i.e. the correct scores (Fig. 1A) in the V-only block. The correct scores varied both across words and subjects from perfect (i.e. 18 correct responses to 18 presentations, e.g. for the word ‘vijf’ by subject S2), to around chance level (0.1, e.g. a score of 0 correct responses for 18 word presentations for the word ‘telde’ presented to subject S8). Notably, some words were easily correctly identified by almost all subjects (e.g. ‘Mark’), while others were near-never identified (‘telde’) by anyone. Similarly, some subjects were perfect lip-readers with correct scores for all words near 1.0 (e.g. subject S14), while subject S13, as an extreme case, could hardly identify any words via lipreading. As the realizations of the visual correct scores were quite noisy (as apparent in the jittery pattern in Fig. 1A), the estimates for the mean correct scores for each word and subject separately (not shown here) were quite uncertain. We therefore determined the visual lipreading recognition rates for words, ρV,w,, and for each subject, ρV,s by fitting the following function:

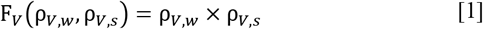

**Fig. 1.**
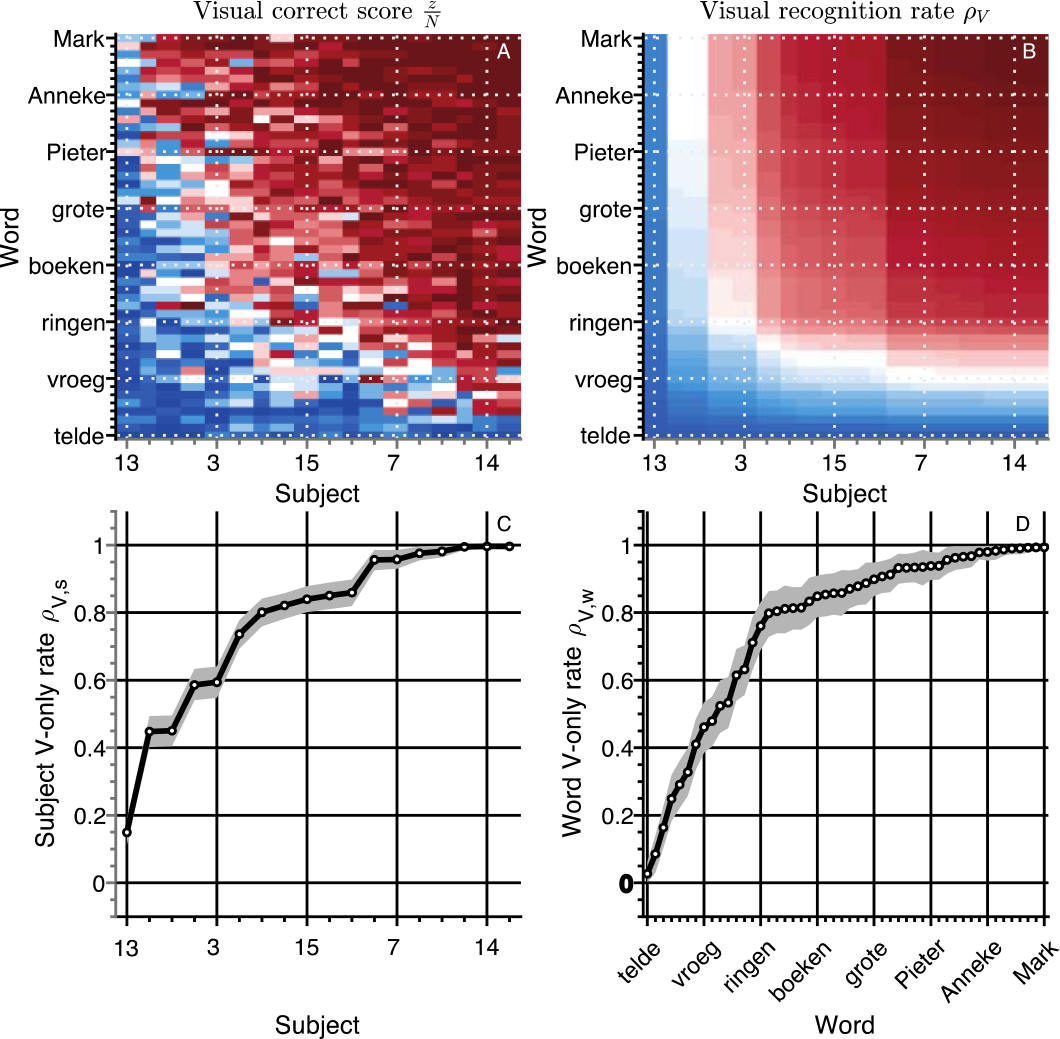
Lip reading. **A)** Visual recognition scores. The correct score (number of correct responses divided by the number of presentations) is shown separately for every word and subject (900 entries) for the V-only condition. The correct scores and rates have been ordered by the recognition rates of subjects on the abscissa, and of words on the ordinate from low-left to high-right. Color codes for correct score or rate, from dark blue = 0 to white = 0.5 to dark red = 1. **B)** Visual recognition rates (Eqn. 1). Same layout as in A. V-only speech recognition rates for **C)** subjects and **D)** words. Rates were ordered from low-left to high-right. Open circles indicate the mean of the estimated rate. Gray patch indicates the 95% Highest Density Interval (HDI).

to all correct and incorrect responses from all V-only trials (see Methods for details). This yields 18 visual recognition rates for subjects, *ρv*,s, and 50 visual recognition rates for words, *ρV*,*w*. Multiplication of these rates assumes that they were independent, and thus separable from each other. This assumption seems to hold, at least qualitatively, when looking at the correct scores for each word and subject (cf. Fig. 1A and Fig. 1B, see also Methods for a more quantitative approach). This procedure smoothened the recognition rate matrix (Fig. 1B), and decreased variability in the estimates (as expressed by the small 95%-HDI in Fig. 1C/D). This function also reduced the number of variables from 900 (number of subjects multiplied by number of words) to 68 (number of subjects plus number of words). These features enable a more practical comparison to the other, A-only and AV conditions, to be introduced later on.

Moreover, the recognition estimates are in line with the correct-score data. Words were generally easily recognized though lipreading (Fig. 1D, mean *ρV,w* = 0.77), but there was considerable variability in visual recognizability across words: many words were identified easily (e.g. mean *ρv,boten* = 0.99), while others were barely recognizable (e.g. mean *ρV,telde* = 0.03). Also the ability of subjects to lipread was relatively high on average (Fig. 1C, mean *ρV,s* = 0.78). However, there was a considerable range in lipreading ability. The best lip-readers could recognize ~100% of the easily-identified words (mean *ρV,S14* = 1.00), while the worst performer could at best recognize ~15% correctly (mean *ρv,s13* = 0.15). The large variability in visual recognition rates across words and subjects provides a potential way to determine how speech-reading performance affects speech-listening, when both auditory and visual speech-recognition cues are presented synchronously.

### Speech listening

In the A-only block, subjects identified words by listening to the audio recordings of sentences (without visual feedback from the lips). A stationary masking noise was played at 65 dB SPL, while the sentences were played at an SNR of −21, −16, −13, −10 or −5 dB. The average word recognition rate was ~50% across all SNRs and subjects (Fig. 2A-E). Overall listening performance for SNRs lower than −10 dB was worse than lipreading performance (cf. amount of red in Fig. 1A vs. blue in Fig. 2A-E). In contrast to lipreading, listening performance was quite similar across subjects (Fig. 2A-E). This small variability across listeners might be expected, as all listeners were normal-hearing, and were therefore likely to understand the speech equally well.

**Fig. 2.**
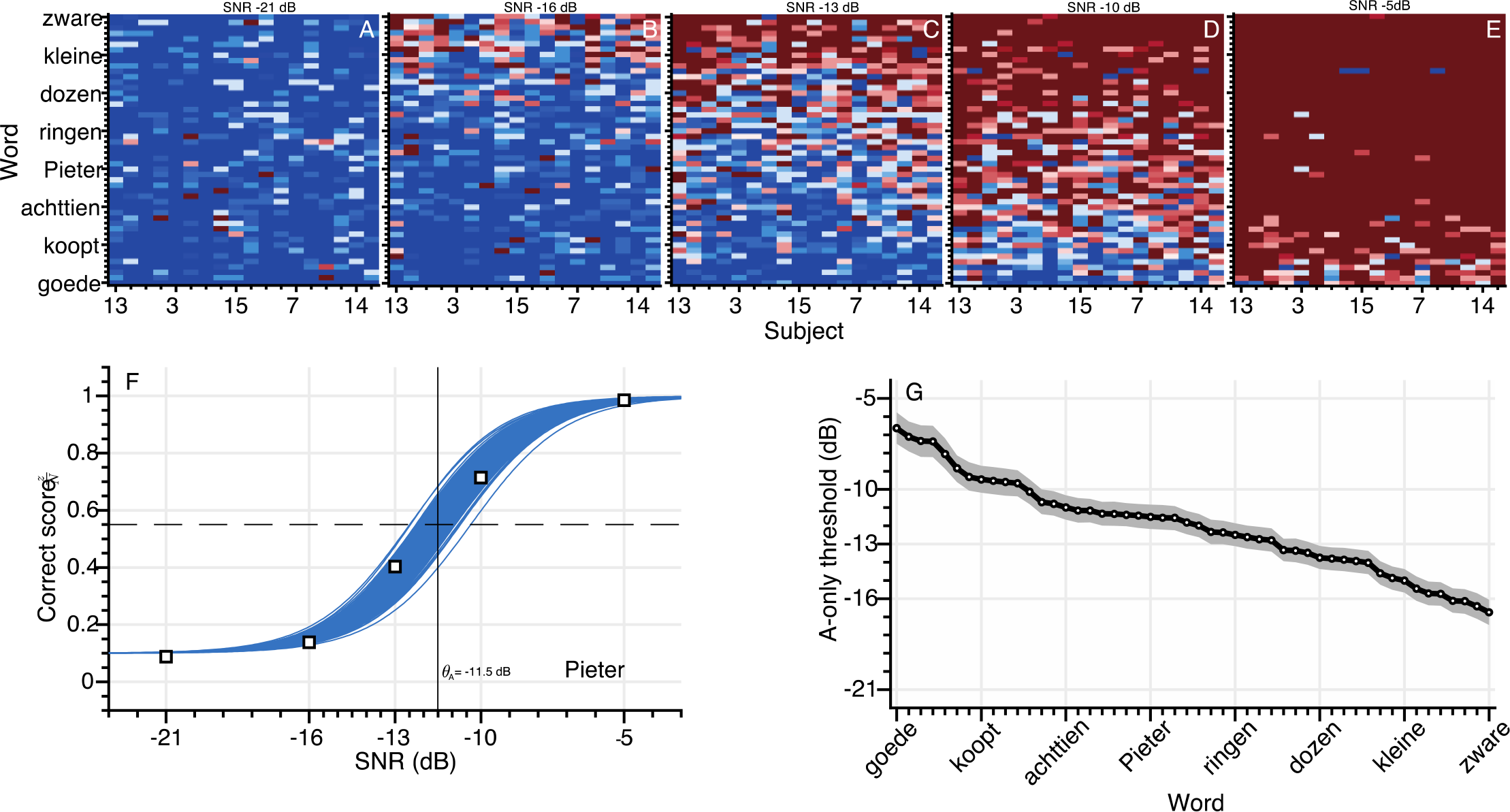
Speech listening. Auditory word-recognition scores. **A-E**) The correct score (number of correct responses divided by the number of presentations) is shown separately for every word and subject (900 entries) for each of the SNRs of −21, −16, −13, −10 and −5 dB. The correct scores have been ordered by the average V-only scores of subjects on the abscissa, and A-only scores of words on the ordinate. Color codes for correct score, from blue = 0 to white = 0.5 to red = 1 (see color bar in Fig. 1). **F)** Correct scores and psychometric fit for the word ‘Pieter’ as a function of SNR, averaged across all subjects. Open squares indicate the measured correct scores. Blue shading denotes credible fits (see Methods). Vertical bold grey line indicates the average of likely recognition thresholds. **G)** A-only speech recognition thresholds, ordered from high-left to low-right. Note that a lower threshold indicates better performance. Open circles indicate means of the estimated thresholds, gray patch indicates the 95% HDI.

Typically, SNR had a strong influence on the ability to recognize the words through listening (Fig. 2A to 2E, from low to high SNR, the correct scores improve from almost 0 to near perfect). To quantify this, we estimated the SNR for which the recognition rate was 50%, i.e. the auditory speech-recognition threshold, *θ*_*A*_, by fitting the parameters of a logistic psychometric function *F*_*A*_ for every word (with a parametrization as mentioned in [31]):

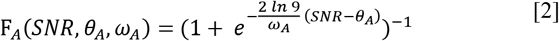

with *ωA* the auditory recognition width from 10 to 90% performance (in dB). The width (conversely, the slope) of the psychometric curve, *ωA*, did not vary substantially across words or subjects. Therefore, only one value was estimated, which was on average 7.1 dB, 95% HDI: 6.8 - 7.4 dB. As the correct scores did not vary appreciably across subjects, we pooled over subjects, to obtain 50 auditory recognition thresholds, one for each word (see also Methods). To exemplify this, we take a look at the word ‘Pieter’ (Fig. 2F). This word was easily recognized by all subjects at the SNR of −5 dB, leading to a 100% recognition score. In contrast, “Pieter” was almost impossible to identify at the lowest SNR of −21 dB, when subjects identified the word presented in 10% of the cases (chance-level). By fitting a psychometric curve through the data, we obtained a speech-listening threshold for this word at −11.5 dB (Fig. 2F, vertical grey bold line). Auditory speech-recognition thresholds for each word (Fig. 2G) varied over a 10-dB range, from the best-recognizable word (mean *θ*_*A*_,_*zware*_ = −16.7 dB) to the hardest-to-recognize word (mean *θ*_*A,goede*_ = −6.6 dB), with an average threshold of −12.1 dB.

### Audiovisual speech recognition

In the AV-condition, subjects identified words by listening to and by lipreading the audiovisual recordings of sentences. The noise was played at 65 dB, while the sentences were played at an SNR of −21, −16, −13, −10, or −5 dB. The presentation of congruent visual feedback clearly aided recognition performance, as the correct scores (Fig. 3A-E) were higher than for the A-only condition (cf. Fig. 2A-E). Also, in contrast to the speech listening scores (cf. Fig. 2A-E) and more in line with lipreading performance (Fig. 1A-B), the AV scores not only varied over words, but also across subjects (which is visible in the pattern of correct scores in Fig. 3A).

**Fig. 3.**
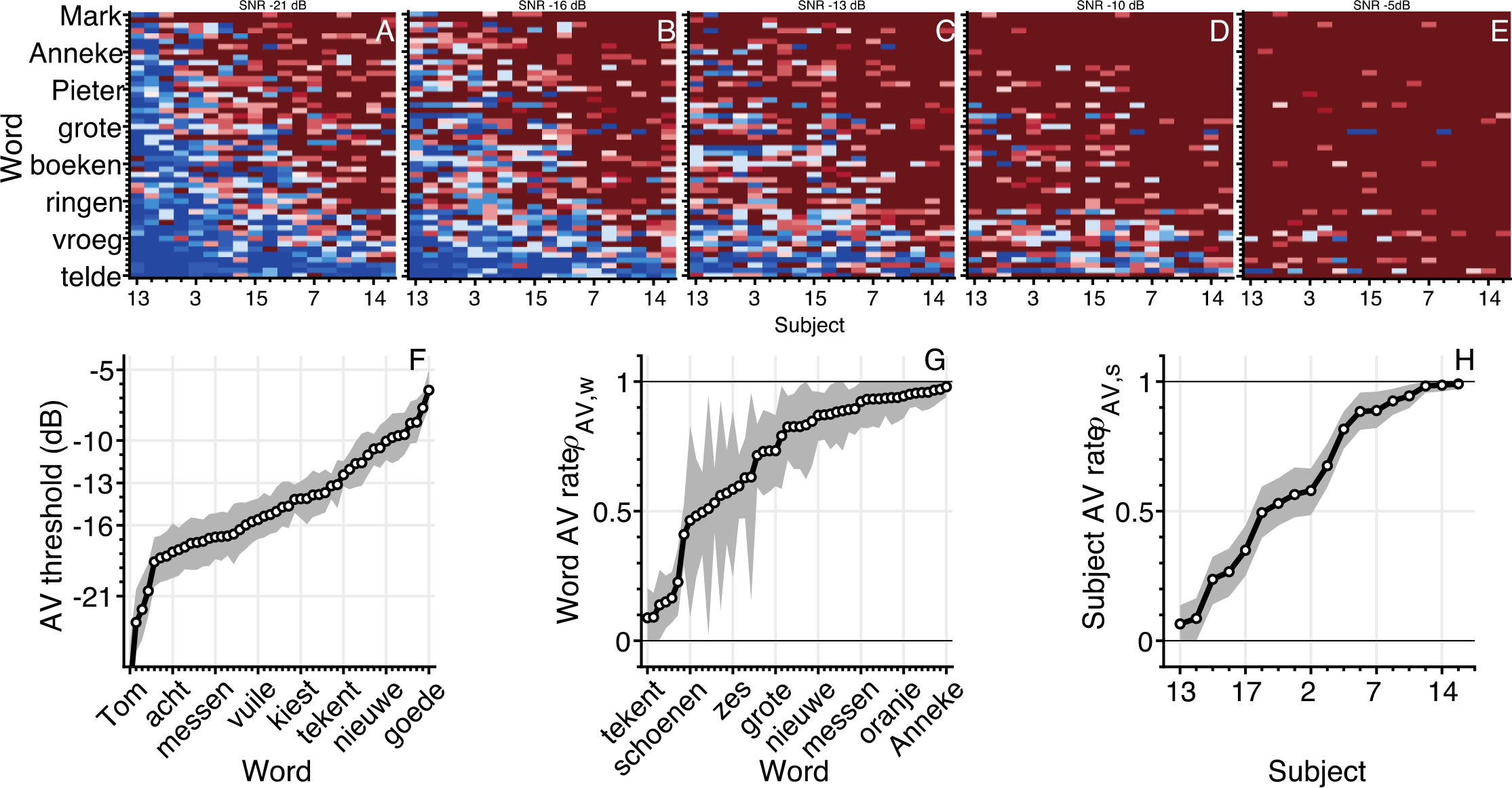
Audiovisual speech recognition. Audiovisual word-recognition scores. **A-E**) The correct score (number of correct responses divided by the number of presentations) is shown separately for every word and subject (900 entries) for each of the SNRs of −21, −16, −13, −10 and −5 dB. The correct scores have been ordered by the average V-only scores of subjects on the abscissa, and of words on the ordinate. Color codes for correct score, from blue = 0 to white = 0.5 to red = 1 (see color bar in Fig. 1). **F)** AV speech-recognition thresholds, **G,H)** AV recognition rates for words and subjects, ordered from low-left to high-right. Note that a lower threshold indicates better performance. Open circles indicate means of the estimated thresholds, gray patch indicates the 95% HDI.

We quantified AV performance by a fitting a function *F*_*AV*_ that combines the characteristics of the eqns. 1 and 2 for the unimodal performances:

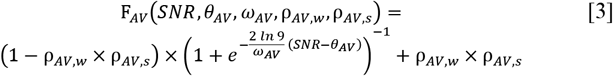

with the audiovisual recognition threshold, *θ*AV describing the logistic SNR dependence, and two audiovisual recognition rates ρAV,w and ρ AV,s, defining the minimum performance level in the AV condition (i.e. for SNR = −∞) for words and subjects, respectively. Like for the A-only condition, one value of the width was estimated for all subjects and words (this width was on average 10.5 dB, 95% HDI: 9.5 - 11.4 dB). The audiovisual speech thresholds were determined for words alone (Fig. 3F), in line with the auditory speech thresholds. The thresholds varied over a ~21 dB range (from mean *θ*_*A, Tom*_ = −27.6 dB to mean *θ*_*A,goede*_ = −6.4 dB), with an average threshold of −14.7 dB. The subjects’ AV recognition rates (Fig. 3G) varied from almost negligible (chance) to near-perfect (from mean *ρAV,S13* = 0.07 to mean *ρAV,S14* = 0.99), with an average rate around 0.63. The AV recognition rates for words (Fig. 3H) varied over a similar range (from mean *ρ*_*AV,tekent*_ = 0.09 to mean *ρ*_*AV,Anneke*_ = 0.98), with an average rate around 0.71. There was considerable uncertainty in the estimation of the word AV rates (e.g., the widest 95%-HDI = 0.02-0.95 for the word ‘Tom’), but in general the 95% HDIs for all other parameters were narrow.

### Audiovisual enhancement

The audiovisual parameters from eqn. 3 are basic descriptors for the audiovisual performance, from which we can derive the audiovisual enhancement by comparing the results to the unimodal parameters from eqns. 1 and 2. For the audiovisual threshold, the comparison to the auditory threshold indicates how much the SNR can decrease when the visual modality is added, without affecting performance. The change in threshold, *Δθ*_*AV*_, relative to the auditory threshold, was thus estimated by rewriting *θ*_*AV*_ in eqn. 3 as:

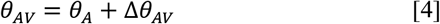

Typically, the audiovisual recognition thresholds were lower (i.e. better) than the auditory recognition thresholds (Fig. 4A), by on average −1.3 dB. This means that the threshold is typically reached at lower SNRs when people speech-read and listen at the same time. The threshold for 35 words improved in the AV condition (95%-HDI lay below 0 dB), while for 15 words there was no difference (95%-HDI included 0 dB). Similarly, the minimum performance level in the AV condition is given by multiplying the recognition rates for words and subjects: *ρ*_*AV,W*_ × *ρ*_*AV,S*_. This measure quantifies the performance level in the absence of an auditory signal (i.e. when the SNR approaches −∞). In case there really is no auditory signal, one might expect that the minimum audiovisual performance level, given by the rates, would equal the visual performance rate. This, of course, only holds if the stimulus parameters fully determine the subject’s performance levels, and if non-stimulus factors, such as task or block design, are irrelevant. We tested this prediction by determining the difference in audiovisual and visual rates for words and subjects:

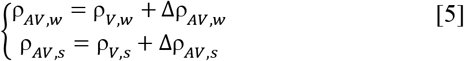

**Fig. 4.**
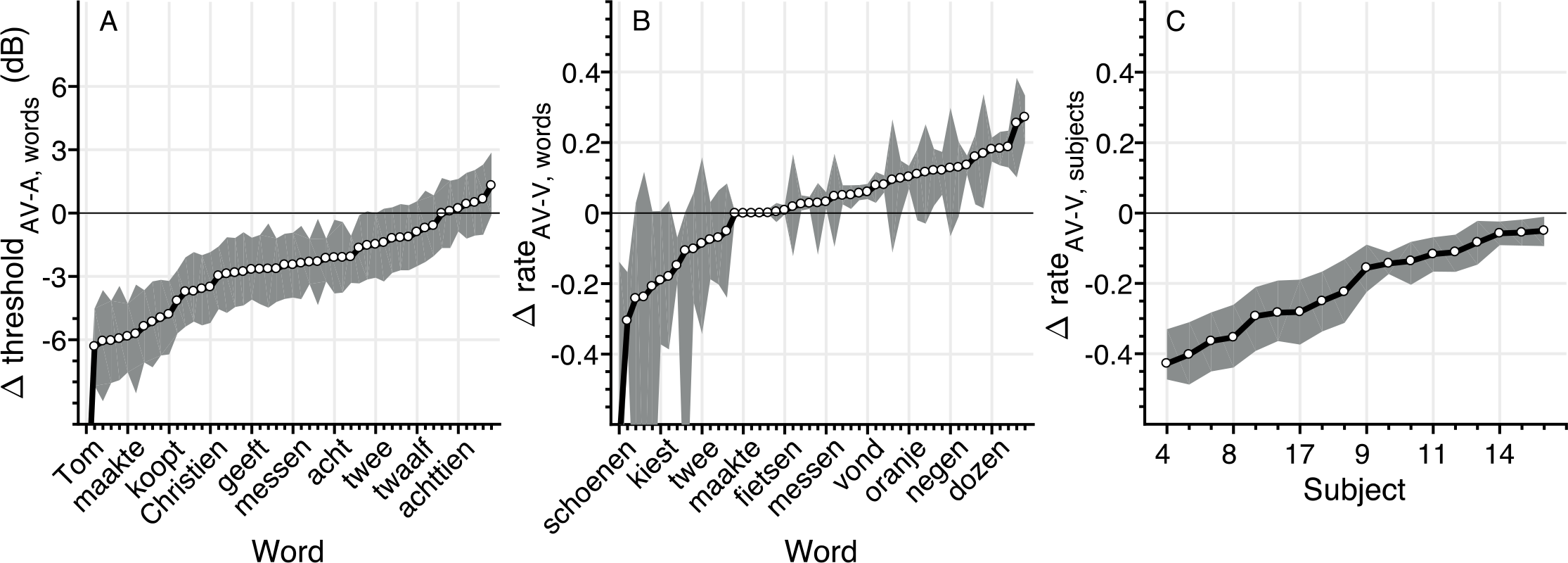
Audiovisual speech recognition. Audiovisual word-recognition scores. **A-E**) The correct score (number of correct responses divided by the number of presentations) is shown separately for every word and subject (900 entries) for each of the SNRs of −21, −16, −13, −10 and −5 dB. The correct scores have been ordered by the average V-only scores of subjects on the abscissa, and of words on the ordinate. Color codes for correct score, from blue = 0 to white = 0.5 to red = 1 (see color bar in Fig. 1). **F)** AV speech-recognition thresholds, **G,H)** AV recognition rates for words and subjects, ordered from low-left to high-right. Note that a lower threshold indicates better performance. Open circles indicate means of the estimated thresholds, gray patch indicates the 95% HDI.

On average, there was no difference in recognition rates for words (Fig. 4B), as the difference values scattered around 0 for most words. In contrast, the subjects’ ability to lipread in the AV condition (as reflected by the subjects’ recognition rate) was poorer than in the V-only condition (Fig. 4C). The rates for all subjects dropped (mean Δρ = −0.2, all 95% HDI < 0). This indicates that, on average, audiovisual performance dropped below the V-only performance scores, when poor auditory SNRs caused speech-listening to deteriorate completely.

As these last points are important, we will restate them. First, the AV threshold is lowered, making it easier to recognize words at a given SNR. This effectively yields an audiovisual enhancement to speech listening (Fig. 4A). Second, words are recognized through lipreading at equal levels in both V-only and AV conditions (Fig. 4B). Third, somewhat surprisingly, the lipreading ability of subjects is impoverished in the AV condition (Fig. 4C). This suggests that task constraints (i.e. being in an AV condition vs. in a V-only condition) have a significant influence on speech recognition performance, even when stimulus parameters are equivalent (i.e. only a visual, no auditory signal).

### Probability summation

Next, we qualitatively compared the AV condition with a model in which audiovisual integration is merely a result of statistical summation rather than of true neural integration. Finding an improved performance (i.e. better speech recognition) in the AV condition is not automatic evidence that the brain integrates the auditory and visual inputs. Indeed, having both modalities available, rather than one, automatically increases the probability of stimulus recognition. In a model of probability summation, participants recognize a word from either the A-only or the V-only condition, which are considered independent processing channels. The probability of word recognition in the presence of the two independent, non-interacting, modalities is given by:

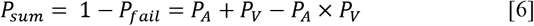

where *P*_*sum*_ is the probability to successfully recognize a word according to the summation model, *P*_*A*_ is the probability to recognize a word in the A-only condition, and *P*_*V*_ is the probability of recognizing a word in the V-only condition. Both *P*_*A*_ and *P*_*V*_ were estimated from eqns. 1 and 2, but were not free parameters of the probability summation model. In order to test how well this model performs for various unimodal stimulus strengths, we split the data in four groups (Fig. 5), as a first, simple approximation, consisting of poor or good lipreading abilities (recognition rate below or above the mean, *ρ*_*V,W*_ = 0.77, respectively) and poor or good auditory thresholds (threshold above or below the mean, *θ*_*A*_ = −12.1dB, respectively). Note that there is a weak correlation between the speech-listening threshold and lipreading recognition rate at the word level; r = −0.39, 95%-HDI = −0.63 to −0.15, so that each group contains a slightly different number of subject-word combinations.

Despite the differences in unimodal performance, the best-fit performance curves (according to eqn. 1-3) for each of those groups followed a similar pattern. Auditory performance (Fig. 5 – blue) degrades as the signal-to-noise ratio decreases; degradation is worse for the words with poor auditory thresholds (Fig. 5A,C). Visual performance (Fig. 5 – red) is better than auditory performance for a larger range of SNRs if the visual word recognition rate is better (Fig. 5A,B). Notably, for all groups, audiovisual performance (Fig. 5 – green) is never worse than auditory performance; a clear audiovisual enhancement relative to auditory performance alone is present for a large range of SNRs. While audiovisual performance is typically also better than visual performance, at very low acoustic SNRs, the multisensory performance tends to be worse than lipreading performance (Fig. 5, the green curves drop below the red lines).

**Fig. 5.**
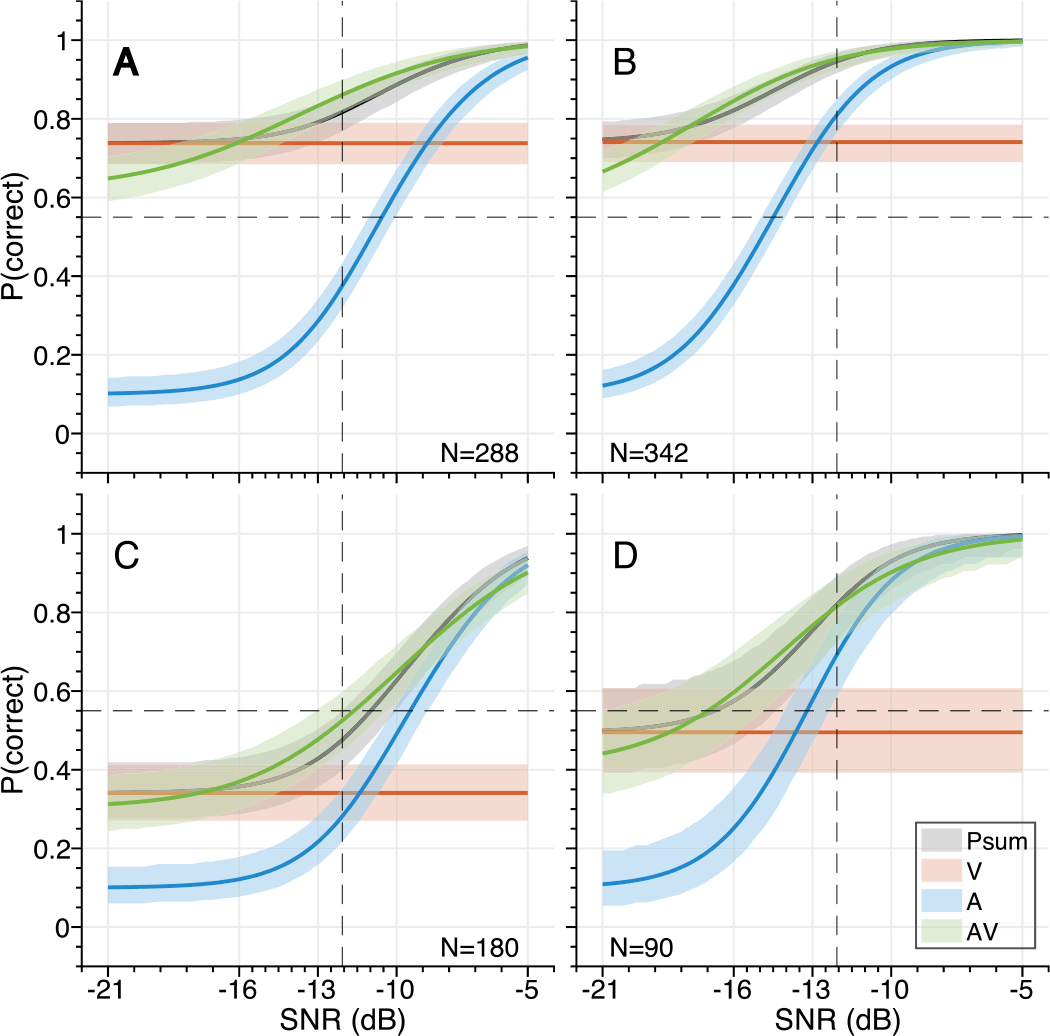
Audiovisual speech recognition varies with unimodal saliency. Psychometric curves were determined (eqn. 1-3) from all data divided across 4 groups differing in unimodal performances: visual recognition rate **A,B)** larger than 0.77 and **C,D)** smaller than 0.77; and an auditory threshold **A,C)** larger than and **B,D)** smaller than −12.1dB. Blue – A-only; red – V-only; green – AV; grey – probability summation. Dashed lines depict the mean (A and V), patches denote the 95% HDI interval. N is the number of subject-word combinations for each group.

Notably, the benchmark probability summation model can describe the audiovisual data quite well, at least qualitatively (Fig. 5 – grey). This model exhibits unimodal-like performance whenever either unimodal recognition abilities vastly outperforms the other, and shows maximum enhancement when the visual and auditory performances are equal. As the audiovisual performance was well described by probability summation, we did not attempt to fit other models exhibiting enhancements [27,29].

### Inverse effectiveness – noise level

To test whether the multisensory data adhered to the principle of inverse effectiveness, we first determined the influence of SNR, as a measure of auditory stimulus intensity, on the magnitude of the audiovisual enhancement. For this purpose, we determined the audiovisual enhancement as the difference between the average best-fit audiovisual and auditory curves (Fig. 5, green and blue). The shape of audiovisual enhancement is largely similar across the four groups. (Fig. 6, blue), and indicates 1) that auditory recognition performance improves by adding the visual information especially for low SNRs, and 2) the highest enhancement occurs at high to intermediate noise levels (SNR between −13 and −20 dB). For lower SNRs, enhancement decreases slightly, so in the strict sense, the principle of inverse effectiveness does not seem to hold for this data set [27].

**Fig. 6.**
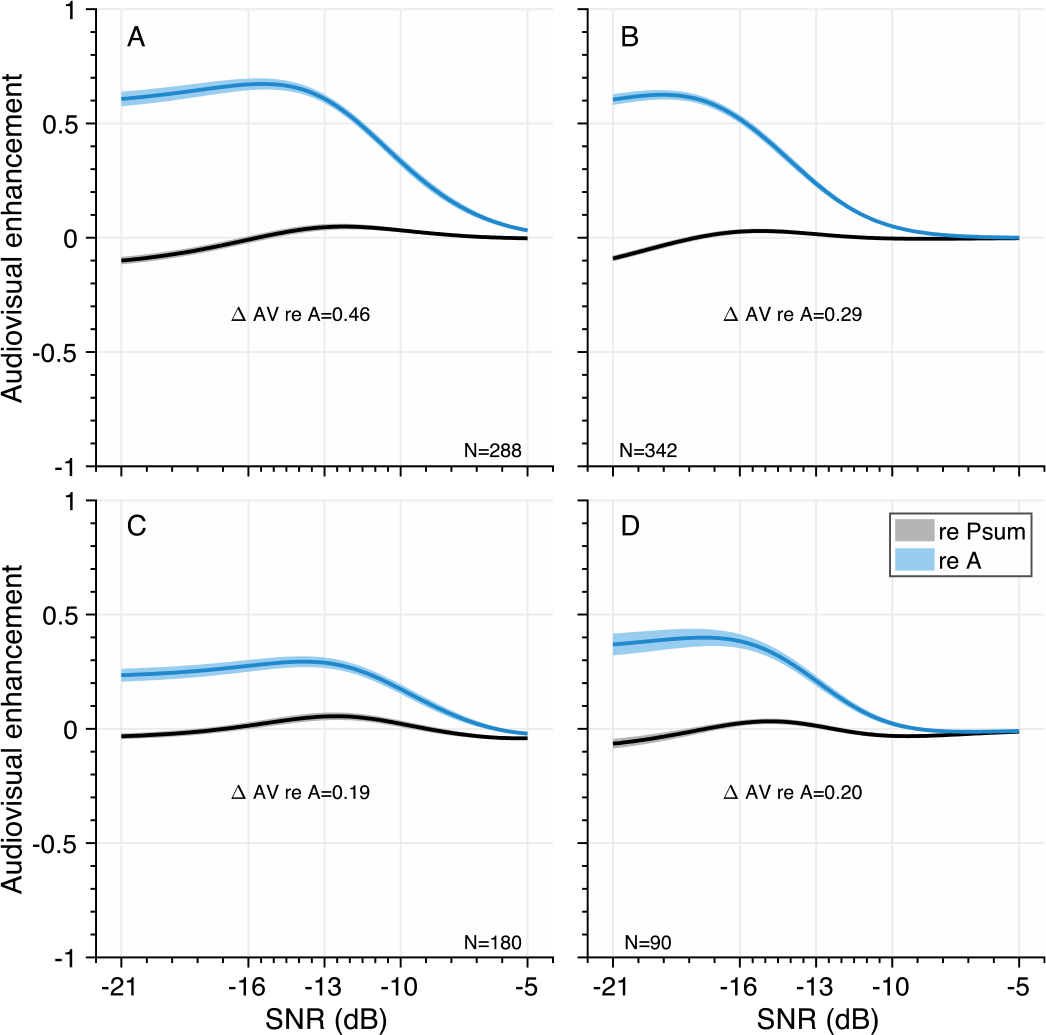
Audiovisual enhancement as a function of SNR. **A-D)** The average audiovisual enhancement, expressed as probability correct, as a function of SNR, compared to speech-listening only (blue) and the probability summation model (black).

We can also express the audiovisual enhancement relative to the benchmark model of statistical summation. For all 4 groups, the probability summation model resembles AV speech recognition quite well (Fig. 6; black lines close to 0). However, there is a slight enhancement over the model at intermediate SNRs (maximum enhancements of 0.03 to 0.06 at SNRs from −12.2 to −15.2 dB), and a slight deterioration at the lowest SNRs (maximum deterioration of −0.04 to −0.10 at an SNR of −21 dB).

### Inverse effectiveness – word saliency and subject performance

Finally, we tested whether multisensory enhancement correlates negatively with unisensory responsiveness (i.e. A-only thresholds, V-only word and subject recognition rates; rather than stimulus intensity, i.e. SNR), as predicted by the principle of inverse effectiveness. To that end, we determined the multisensory enhancement as the difference in recognition probability between the audiovisual and either the auditory, *E*_*AV-A*_, or visual, *E*_*AV-V*_, stimulus, averaged across SNR, for every word and subject. The slope of the relationship between multisensory enhancement and auditory thresholds or visual recognition rates, respectively, was determined through multiple linear regression analysis:

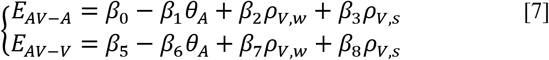

with *β*_*1*_ the parameter of interest to infer effectiveness of the auditory response, and *β*_*7*_ and *β*_*8*_ of the visual response for words and subjects. The other parameters are included to account for confounds such as the effect of the other modality (e.g. the audiovisual enhancement over the auditory response will be negligible if the visual response is minimal). These parameters are an offset to the intercept and reflect the type of integration as shown by the audiovisual data (i.e. super-additive, additive, sub-additive). Note that for the auditory thresholds, the signs are inverted. This ensures that a negative slope would actually indicate inverse effectiveness, even though higher thresholds indicate a worse response.

The audiovisual enhancement over the auditory response *(E*_*AV-A*_, Fig. 7A) is larger for words with higher auditory thresholds, with an effectiveness slope *β*_1_ = −0.036 (95%-HDI: −0.038 to −0.033). The negative slope suggests that the auditory response to each word is inversely effective in driving the multisensory response. The magnitude of the enhancement over the auditory response increases when a word can be more easily recognized through lip-reading (i.e. high visual word recognition rate, dark filled dots). This is in line with the observation that the multisensory data follow probability summation quite well, reflecting an additive type of integration (Fig. 5 and 6). Importantly, the observed inverse effectiveness is not an artefact due to a ceiling effect, as the auditory response allowed for a larger performance benefit (Fig. 7A, dotted line).

**Fig. 7.**
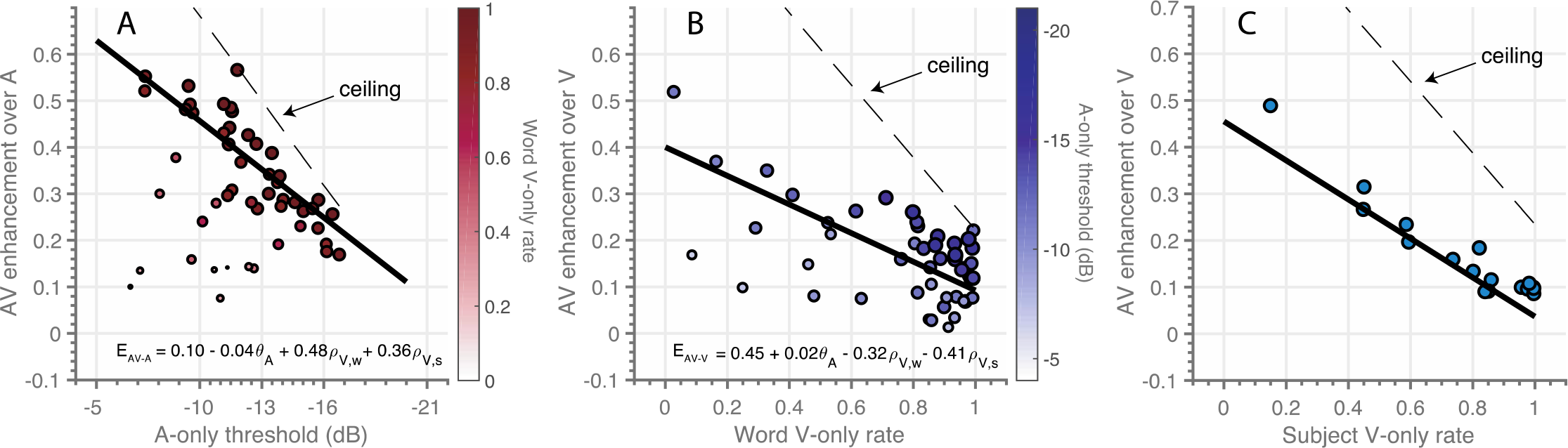
Inverse effectiveness. The audiovisual enhancement over unisensory responses (as defined in the text) as a function of **A)** auditory threshold, **B)** visual word recognition rate, **C)** visual subject recognition rate. Note that the x-axis is inverted in A). Black lines indicate the best-fit multiple regression line for the independent variable of interest, with an intercept determined by the median of the other variables. Dashed lines indicate the maximum enhancement possible (1.0 - unisensory performance, for the given condition). Dot size and color in A) and B) denote the cross-sensory performance level as indicated by the color bars.

Multisensory enhancement over the visual response follows the same principles. Words with a low visual recognition rate were more effective at improving the AV response (Fig. 7B), with an effectiveness slope *β*_*7*_ = −0.32 (95%-HDI: −0.34 to −0.29). Notably, even across subjects, the poorer lipreaders benefit more from audiovisual presentation than excellent lipreaders (Fig. 7C), with an effectiveness slope *β_8_ =* −0.41 (95%-HDI: −0.44 to −0.38).

## Discussion

### Overview

This paper reports the occurrence of inverse effectiveness on the recognizability – visually or auditory - of individual words. We determined how well words presented in sentences can be recognized by normal-hearing subjects through listening and/or lipreading under noisy listening conditions. In line with previous research [4–7], we found that lipreading improves speech recognition by listening alone (Fig. 4A, Fig. 5). However, we also observed that audiovisual performance levels fall below lipreading performance for the lowest SNR (Fig. 4C, Fig. 5). Furthermore, we found that the largest improvements were typically found at intermediate SNRs, rather than at the lowest SNRs, which is line with previous research [27]. Although this discredits the principle of inverse effectiveness in audiovisual speech perception, we did observe inverse effectiveness across individual words and subjects (Fig. 7): the data show that the benefit of adding cross-modal information increased when a word was poorly heard (Fig. 7A), when a word was poorly seen (Fig. 7B), or when the subject was a poor lipreader (Fig. 7C). To our knowledge, this has not been shown so far.

### Performance in lipreading

Our data demonstrate considerable variability in lipreading performance (Fig. 1), which has been reported and discussed earlier in the literature [32]. The average performance levels from the current study are relatively high, especially considering that the normal-hearing subjects were not specifically trained to lipread. This is consistent with earlier findings on word and sentence recognition tasks [32], although more recent papers have reported lower values [11,27,29]. Bernstein et al. [32] questioned whether it was actually possible for normal-hearing subjects to have good lipreading abilities. One possible explanation for the high lipreading performance might be the use of the closed-set speech-recognition task (i.e. a limited set of words used in the behavioral task). However, one would then also expect to observe a familiarization or training effect over sessions, which did not occur (data not shown).

### Performance in speech listening

The auditory scores varied mainly across words; subjects could all recognize words through listening at an almost equal performance level (Fig. 2). Since all participants had normal hearing, and could therefore be expected to understand speech equally well, the limited variability between subjects corroborated that expectation. The analysis of speech recognition performance in the auditory-only condition revealed speech reception thresholds of −12.1 dB, which is lower than the threshold of −8.4 dB obtained from the original version of the Dutch Matrix test [30]. Models for audiovisual enhancement

The behavioral improvement of audiovisual speech perception can be modeled in various ways. Typically, AV data are compared to the benchmark probability-summation model, in which the auditory and visual channels are considered independent, without true multisensory neural interactions. This model was the only model considered in this study (Eqn. 6, Figs. 5 and 6), as it matched the data closely (Figs. 5 and 6).

Rouger and colleagues [29] found that an alternative, optimal-integration model could better describe their data. In their model, spectral-temporal audiovisual cues merge across modalities to optimize the amount of information required for word recognition. Our audiovisual data in poor lipreading conditions (i.e. visual recognition rate for a word is lower than 0.5) compares quite well to the speech-recognition abilities of the normal-hearing subjects of Rouger et al. in the presence of a masking noise (cf. [29] - their Fig. 3D).

A third model was proposed by Ma and colleagues, in which words were regarded as points in a multidimensional space, and word recognition becomes a probabilistic inference process [27]. This Bayesian model assumes that certain words occur more frequently than other words (and are more easily recognized), and it uses this pre-knowledge (i.e. priors) to explain the recognition scores for all words.

It is hard to reconcile any of the three models with our observation that in low-SNR conditions, multisensory speech recognition is actually degraded compared to unimodal lipreading without accounting for non-stimulus factors affecting audiovisual speech recognition (Figs. 4C and 5). The aforementioned models do not include a mechanism for divided attention between the two modalities [33,34]. In such a scheme, the two separate information streams could actually lead to impaired performance in conditions in which either of the two signals may be ambiguous or weak. Thus, even though lipreading might provide sufficient information to recognize words, people are not able to divert their attention away from the auditory stream, despite the absence of a potential signal in that information stream.

### Inverse effectiveness

We tested whether the principle of inverse effectiveness also holds in audiovisual speech recognition by: i) modulating the acoustic signals re background noise, ii) by investigating each subject’s lipreading ability, and iii) by comparing to auditory and/or visual recognizability of words. First, in line with several laboratory studies of multisensory integration using simple sensory stimuli (e.g. white noise bursts and LED flashes) [16-23,25,26], a lower auditory SNR typically induced stronger multisensory enhancement. However, here we report that for the lowest SNRs (−21 dB) the enhancement saturated, or even slightly dropped (Fig. 6). This differs quantitatively with the data from Ma et al. [27], who found a significant enhancement drop for low and high SNRs. Qualitatively, however, both studies provided evidence for the lack of inverse effectiveness when the acoustic cues were impoverished, due to reduced signal levels in the presence of noise. Notably, Bayesian modelling of audiovisual enhancement in the study by Ma et al. suggested that the largest enhancement shifted to lower SNRs with decreasing vocabulary size. As the vocabulary size in the current experiment was limited to only 50 words, the model by Ma et al. would also predict the largest enhancement at the lowest SNRs.

Secondly, evidence for inverse effectiveness can be found for individual lipreading abilities; worse lipreaders benefitted more from the additional auditory information for the audiovisually presented sentences (Fig. 7C). Due to ceiling effects, this might appear trivial, as the best lipreaders cannot improve their performance further by similar amounts as the poorest lipreaders (see [35,36]).

Finally, inverse effectiveness also plays a role at word-level performance, both for vision and for hearing: the hardest to-recognize words underwent the strongest audiovisual enhancements relative to the unimodal condition (Fig. 7). As such, sensory conditions and perceptual responses seemed to be more in line with basic multisensory integration results from earlier studies using simple noise bursts and LED flashes and even for studies using complex, spectro-temporally modulating stimuli [24].

### Matrix test

The audiovisual speech material is based on an existing auditory-only matrix sentence test for Dutch native speakers [30,37]. It is not immediately clear whether the observed results hold specifically for the Dutch language, or whether it is immaterial for which language this test has been developed. Numerous audiovisual speech recognition tests have been developed for the English language [4,9,11,13,27,38]. Exceptions are studies for native French [29,39] and Dutch speakers [40]. Detailed comparisons are difficult also because the stimuli (monosyllables vs words vs sentences) and the subject populations (normal-hearing vs hearing-impaired) differ. The use of a standardized test, such as the Matrix test, might facilitate comparisons, especially between normal-hearing and hearing-impaired listeners, since the Matrix test is also well-suited to test the hearing-impaired. Comparisons across languages might still be difficult, as, even though an auditory Matrix test is available in many languages [30,41-43], the words may vary in their spectro-temporal properties and thresholds between languages.

### Conclusion

To conclude, lipreading enhances speech recognition (in line with earlier studies); this visual enhancement, however, is affected by the acoustic properties of the audiovisual scene. Visual enhancement for words that are easily recognized by vision alone is impoverished in high acoustic noise conditions. Audiovisual enhancements were highest for intermediate signal-to-noise ratios. Inverse effectiveness holds for words and subjects, for which the poorest visually/auditory-recognizable words underwent the strongest cross-modal enhancements.

## Materials and Methods

### Participants

Eighteen native Dutch-speaking adults (mean age = 26 years, range = 21-40) participated in this study. All gave their informed consent. They were screened for normal-hearing (within 20 dB HL range 0.5 - 8 kHz), and had normal or corrected-to-normal vision. The experiments were carried out in accordance with the relevant institutional and national regulations and with the World Medical Association Helsinki Declaration as revised in March 2017 (Declaration^1^). The experiments were approved by the Ethics Committee of Arnhem-Nijmegen (project number NL24364.091.08, October 18, 2011).

### Audiovisual material

The speech material was based on the Dutch version of the speech-in-noise matrix test developed by Houben and colleagues [30] in analogy to a Swedish test [41]. In general, a matrix test uses complete sentences that are composed from a fixed matrix of words (Table 1). All created sentences shared the same grammatical structure (name, verb, numeral, adjective, object), but were semantically unpredictable. In principle, a set of 10^5^ different sentences could be created. Therefore, the test suffered little from potential training confounds when participants were tested multiple times. Houben et al. [30], ensured that the occurrence of phonemes in their test was similar to standard Dutch. For the audiovisual version of the test reported here, we selected a subset of 180 (155 unique) sentences that were grouped into 9 lists of 20 sentences each. In every list, each of the 50 words from the matrix occurred twice, once in the first ten sentences and once in the second ten sentences.

**Table 1.**
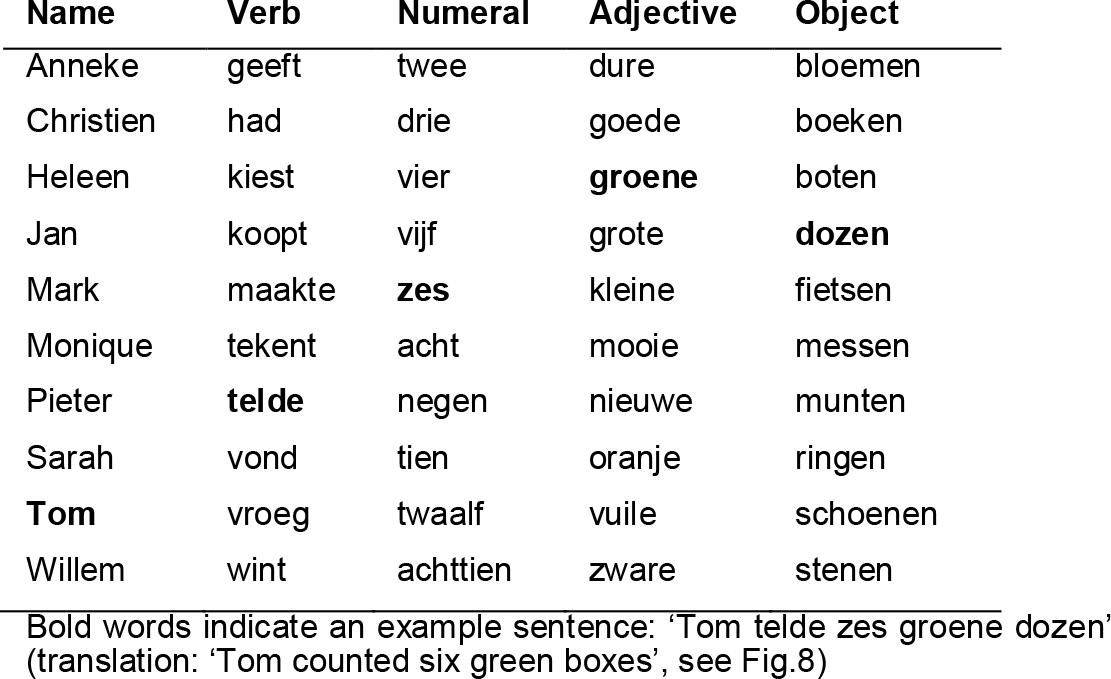
Words of the Dutch matrix test

The audio-video material was recorded in a sound-attenuated, semi-anechoic room, using an Olympus LS-5 audio recorder (24-bit/44.1 kHz sampling rate), and a Canon 60D video camera (1280 × 720, 720p HD at 50 frames per second), respectively. All sentences were spoken by a Dutch female speech therapist. If a sentence was not articulated clearly, or if there was a sudden movement of the face or eyes, the sentence was re-recorded. The audio and video recordings were combined off-line using Final Cut Pro X (Mac App OS X Yosemite), and saved in MPEG-4 format, in H.264 codec.

We applied audiovisual sentence material that was created on the basis of the well-defined audiological Matrix test. With its potential application to a multitude of languages [30,41-43], our study facilitates the comparison of audiovisual speech-in-noise data across languages, as well as between normal-hearing and hearing-impaired listeners.

### Experimental setup

Audiovisual testing was carried out in the same room in which the material had been recorded. Stimulus presentation was controlled by a Dell PC (Dell Inc., Round Rock, TX, USA) running Matlab version 2014b (The Mathworks, Natick, MA, USA). Participants were seated at a table, 1.0 m in front of a PC screen (Dell LCD monitor, model: E2314Hf, Dell Inc., Texas, USA). Sounds were played through an external PC sound card (Babyface, RME, Germany) and presented over one speaker (Control Model Series, model number: Control One, JBL, California, USA) placed 1.0 m in front of the participant, immediately above the screen (30° above the interaural plane). Speaker output was calibrated with an ISO-TECH Sound Level Meter (type SLM 1352P) at the position of the listener’s head, on the basis of the stationary masking noise.

### Stimuli

The stimuli contained digital video recordings of a female speaker reading aloud the sentences in Dutch (Fig. 8). In the auditory-only presentation (A-only), the voice was presented without visual input (i.e. black screen, Fig. 8A,C) with added background acoustic noise (Fig. 8B). In the visual-only presentation (V-only) the video fragments of the female speaker were shown on the screen without an auditory speech signal and noise (Fig. 8D). In the audiovisual presentation (AV), the video was presented with the corresponding auditory signal and the masking noise.

**Fig. 8.**
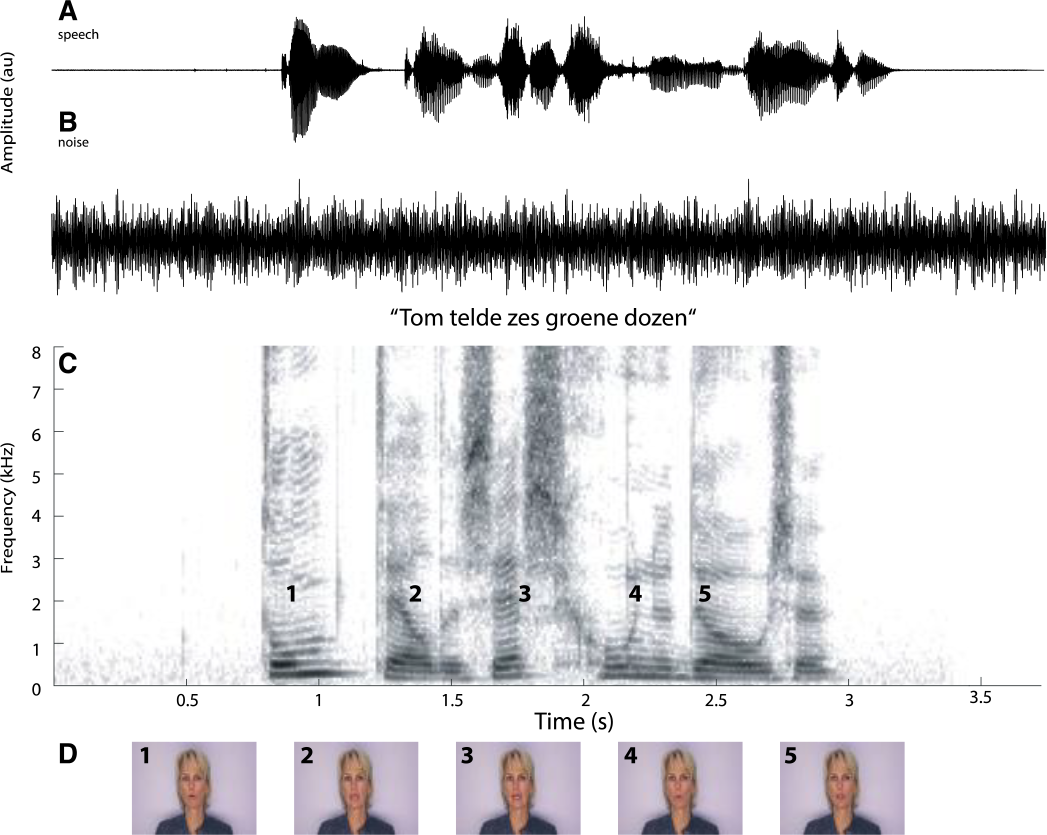
Example stimulus. **A)** amplitude plot of the auditory speech signal. **B)** amplitude plot of the auditory noise. **C)** Spectrogram of the recorded sentence “Tom telde zes groene dozen” (translation: Tom counted six green boxes). Numbers 1-5 denote the occurrence of the video frames shown in D. **D)** Five video frames taken from the video-recorded sentence

The masking noise was created following the procedure reported by Wagener et al. [44]. To that end, the 180 sentences were overlaid by applying a random circular shift. Repeating that procedure five times resulted in a stationary masking noise with the same spectral characteristics as the original speech material.

### Paradigm

All participants were tested in a closed-set speech-recognition test in A-only, V-only and AV conditions. Prior to the experiment, all participants familiarized themselves with the matrix of 50 words (10 words for each of the 5 categories, Table 1) and by practicing the task on 10 randomly selected AV sentences. No improvement in speech recognition was observed during the experimental sessions, which indicates that there was no recognition effect of procedural learning.

The masking noise started and ended 500 ms before and after the sentence presentation. The noise onset and offset included 250 ms (sin^2^, cos^2^) ramps. In the A-only and AV conditions, the masking noise was fixed at 65 dB SPL (A-weighted), with the speech sound presented at 44, 49, 52, 55, or 60 dB SPL (A-weighted) to obtain signal-to-noise ratios (SNRs) of −21, −16, −13, −10, and −5 dB, respectively. After presentation of the sentence and the end of the noise, the matrix of 50 words was shown on the screen (Table 1). Participants were instructed to choose one word from each of the 5 categories (10-alternative forced-choice task). Participants initiated the next trial by pressing the mouse-button.

For each of the sensory modalities (A-only, V-only, and AV), participants were tested in separate sessions on different days. In this way, fatigue and repetitive stimulus presentation were avoided. In each session, the nine lists of 20 sentences were presented. In the A-only and AV sessions, each sentence was assigned one of the five SNRs pseudo-randomly (each SNR was presented equally often as the others, i.e. 36 times in each session).

### Data analysis

For every word (w=1:50), subject (s=1:18), SNR (n=1:5) and sensory modality (m=1:3), we determined the correct score, defined as the number of correct responses, z, divided by the number of presentations, N. The correct score is binomially distributed, in which the probability of a success given by:

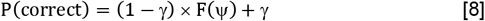

where *F(ψ)* is a function (binomial distribution) that characterizes the recognition performance for the particular stimulus and subject parameters (subject parameters such as SNR and visual recognition rate), described by *ψ*; *γ* is the probability that the subject gives the correct answer, irrespective of the stimulus (the ‘guess rate’). Here, *γ* was set to 10% (0.1), as there were ten word alternatives per category. From the correct scores, we estimated the recognition rates, *ρ* (i.e. how often words were recognized correctly at a given SNR), and the recognition thresholds, *θ* (i.e. the SNR at which words were recognized in 50% of the presentations), as described below.

### Statistical Analysis

Parameter estimation of Eqns. 1-8 was performed using a Bayesian statistical analysis. This analysis requires the definition of priors over the parameters. As a prior for the auditory thresholds, we chose the Gaussian distribution with mean 0 and standard deviation 100, and for the visual recognition rates we took a positive-only beta distribution, for which both shape parameters were set to 1. The audiovisual rate differences (Eqn. 5) were modeled as Gaussian distributions with the rates transformed to probit scale (see e.g. [45] Chapter 9.3). For the multiple linear regression (eqn. 7; [46]), the data was modeled according to a t-distribution. For the priors on the parameters, Gaussian distributions with a mean of 0 and a standard deviation of 2 were chosen, after normalization of the data.

The estimation procedure relied on Markov Chain Monte Carlo (MCMC) techniques. The estimation algorithms were implemented in JAGS [47] through matJAGS [48]. Three MCMC chains of 10,000 samples were generated. The first 10,000 samples were discarded as burn-in. Convergence of the chains was determined visually, by checking that the shrink factor 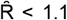 [49,50], and by checking that the effective sample size >1000 [51].

From these samples of the posterior distributions, we determined the mean and the 95% highest density interval (95%-HDI) as a centroid and uncertainty estimate of the parameters, respectively.

## Author Contributions and Notes

LPHVDR, MMVW, AJVO, EAMM and AR designed research, LPHVDR performed research, MMVW wrote software, LPHVDR and MMVW analyzed data; and LPHVDR, MMVW, AR, AJVO and EAMM wrote the paper.

The authors declare no conflict of interest. The funders had no role in study design, data collection and analysis, decision to publish, or preparation of the manuscript.

## Acknowledgments

We thank Günther Windau, Ruurd Lof, Stijn Martens, and Chris-Jan Beerendonck for their valuable technical assistance, speech-therapist Jeanne van der Stappen for providing the audiovisual material, and Eefke Lemmen for editing. We are grateful to Ad Snik for providing valuable comments on earlier versions of this manuscript. This research was supported by EU FP7-PEOPLE-2013-ITN iCARE (grant 407139, AR), EU Horizon 2020 ERC Advanced Grant ORIENT (grant 693400, AJVO), Cochlear Benelux NV (LPHVDR, MMVW), the Radboud University Medical Center (LPHVDR, EAMM), and Radboud University (MMVW).

## Footnotes

1 https://www.wma.net/policies-post/wma-declaration-of-helsinki-ethical-principles-for-medical-research-involving-human-subjects

## References

1. van de Rijt LPH, van Opstal AJ, Mylanus EAM, Straatman LV, Hu HY, Snik AFM, van Wanrooij MM. Temporal Cortex Activation to Audiovisual Speech in Normal-Hearing and Cochlear Implant Users Measured with Functional Near-Infrared Spectroscopy. Front Hum Neurosci. 2016;10: 48. doi:10.3389/fnhum.2016.00048

2. Calvert GA, Campbell R, Brammer MJ. Evidence from functional magnetic resonance imaging of crossmodal binding in the human heteromodal cortex. Curr Biol. 2000;10: 649–57. doi:10.1016/-9822(00)00513-3

3. Stein BE, Meredith MA. The Merging of the Senses. Cambridge, MA, US: The MIT Press.; 1993.

4. Bernstein LE, Auer ET, Takayanagi S. Auditory speech detection in noise enhanced by lipreading. Speech Commun. 2004;44: 5–18. doi:10.1016/j.specom.2004.10.011

5. Grant KW, Seitz PF. The use of visible speech cues for improving auditory detection of spoken sentences. J Acoust Soc Am. 2000;108: 1197–208. doi:10.1121/1.422512

6. Helfer KS. Auditory and auditory-visual perception of clear and conversational speech. J speech, Lang Hear Res. 1997;40: 432–43.

7. Winn MB, Rhone AE, Chatterjee M, Idsardi WJ. The use of auditory and visual context in speech perception by listeners with normal hearing and listeners with cochlear implants. Front Psychol. Frontiers; 2013;4: 824. doi:10.3389/fpsyg.2013.00824

8. MacLeod A, Summerfield Q. Quantifying the contribution of vision to speech perception in noise. Br J Audiol. 1987;21: 131–41.

9. MacLeod A, Summerfield Q. A procedure for measuring auditory and audio-visual speech-reception thresholds for sentences in noise: rationale, evaluation, and recommendations for use. Br J Audiol. 1990;24: 29–43. doi:10.3109/03005369009077840

10. O’Neill JJ. Contributions of the visual components of oral symbols to speech comprehension. J Speech Hear Disord. American Speech-Language-Hearing Association; 1954;19: 429–439. doi:10.1044/jshd.1904.429

11. Ross LA, Saint-Amour D, Leavitt VM, Javitt DC, Foxe JJ. Do you see what I am saying? Exploring visual enhancement of speech comprehension in noisy environments. Cereb Cortex. 2007;17: 1147–53. doi:10.1093/cercor/bhl024

12. Sommers MS, Tye-Murray N, Spehar B. Auditory-visual speech perception and auditory-visual enhancement in normal-hearing younger and older adults. Ear Hear. 2005;26: 263–75. doi:10.1097/00003446-200506000-00003

13. Sumby WH, Pollack I. Visual Contribution to Speech Intelligibility in Noise. J Acoust Soc Am. 1954;26: 212–215. doi:10.1121/1.1907309

14. Tye-Murray N, Sommers MS, Spehar B. Audiovisual integration and lipreading abilities of older adults with normal and impaired hearing. Ear Hear. 2007;28: 656–68. doi:10.1097/AUD.0b013e31812f7185

15. Tye-Murray N, Sommers M, Spehar B, Myerson J, Hale S. Aging, Audiovisual Integration, and the Principle of Inverse Effectiveness. Ear Hear. 2010;31: 1. doi:10.1097/AUD.0b013e3181ddf7ff

16. Corneil BD, van Wanrooij MM, Munoz DP, van Opstal AJ. Auditory-visual interactions subserving goal-directed saccades in a complex scene. J Neurophysiol. 2002;88: 438–54. doi:10.1152/jn.2002.88.1.438

17. van Barneveld DCPBM, van Wanrooij MM. The influence of static eye and head position on the ventriloquist effect. Eur J Neurosci. 2013;37: 1501–10. doi:10.1111/ejn.12176

18. Alais D, Burr D. No direction-specific bimodal facilitation for audiovisual motion detection. Brain Res Cogn Brain Res. 2004;19: 185–94. doi:10.1016/j.cogbrainres.2003.11.011

19. Körding KP, Beierholm U, Ma WJ, Quartz S, Tenenbaum JB, Shams L. Causal inference in multisensory perception. Sporns O, editor. PLoS One. Public Library of Science; 2007;2: e943. doi:10.1371/journal.pone.0000943

20. Wallace MT, Roberson GE, Hairston WD, Stein BE, Vaughan JW, Schirillo JA. Unifying multisensory signals across time and space. Exp brain Res. 2004;158: 252–8. doi:10.1007/s00221-004-1899-9

21. Bell AH, Meredith MA, van Opstal AJ, Munoz DP. Crossmodal integration in the primate superior colliculus underlying the preparation and initiation of saccadic eye movements. J Neurophysiol. 2005;93: 3659–73. doi:10.1152/jn.01214.2004

22. Meredith MA, Stein BE. Spatial factors determine the activity of multisensory neurons in cat superior colliculus. Brain Res. 1986;365: 350–4. doi:10.1016/0006-8993(86)91648-3

23. Wallace MT, Meredith MA, Stein BE. Multisensory integration in the superior colliculus of the alert cat. J Neurophysiol. 1998;80: 1006– 10. doi:10.1152/jn.1998.80.2.1006

24. Bremen P, Massoudi R, van Wanrooij MM, van Opstal AJ. Audio-Visual Integration in a Redundant Target Paradigm: A Comparison between Rhesus Macaque and Man. Front Syst Neurosci. Frontiers; 2017;11: 89. doi:10.3389/fnsys.2017.00089

25. Frens MA, van Opstal AJ, van der Willigen RF. Spatial and temporal factors determine auditory-visual interactions in human saccadic eye movements. Percept Psychophys. 1995;57: 802–16. doi:10.3758/BF03206796

26. van Wanrooij MM, Bell AH, Munoz DP, van Opstal AJ. The effect of spatial-temporal audiovisual disparities on saccades in a complex scene. Exp brain Res. Springer-Verlag; 2009;198: 425–437. doi:10.1007/s00221-009-1815-4

27. Ma WJ, Zhou X, Ross LA, Foxe JJ, Parra LC. Lip-reading aids word recognition most in moderate noise: a Bayesian explanation using high-dimensional feature space. PLoS One. 2009;4: e4638. doi:10.1371/journal.pone.0004638

28. Tye-Mmurray N, Spehar B, Myerson J, Hale S, Sommers M. Lipreading and audiovisual speech recognition across the adult lifespan: Implications for audiovisual integration. Psychol Aging. 2016;31: 380–389. doi:10.1037/pag0000094

29. Rouger J, Lagleyre S, Fraysse B, Deneve S, Deguine O, Barone P. Evidence that cochlear-implanted deaf patients are better multisensory integrators. Proc Natl Acad Sci U S A. 2007;104: 7295–300. doi:10.1073/pnas.0609419104

30. Houben R, Koopman J, Luts H, Wagener KC, van Wieringen A, Verschuure H, et al. Development of a Dutch matrix sentence test to assess speech intelligibility in noise. Int J Audiol. Taylor & Francis; 2014;53: 760–3. doi:10.3109/14992027.2014.920111

31. Kuss M, Jäkel F, Wichmann FA. Bayesian inference for psychometric functions. J Vis. 2005;5: 478–92. doi:10.1167/5.5.8

32. Bernstein LE, Demorest ME, Tucker PE. Speech perception without hearing. Percept Psychophys. 2000;62: 233–52. doi:10.3758/BF03205546

33. Alsius A, Navarra J, Campbell R, Soto-Faraco S. Audiovisual Integration of Speech Falters under High Attention Demands. Curr Biol. Elsevier; 2005;15: 839–843. doi:10.1016/j.cub.2005.03.046

34. Bonnel AM, Hafter ER. Divided attention between simultaneous auditory and visual signals. Percept Psychophys. Springer-Verlag; 1998;60: 179–90. doi:10.3758/BF03206027

35. Stein BE, Stanford TR, Ramachandran R, Perrault TJ, Rowland BA. Challenges in quantifying multisensory integration: alternative criteria, models, and inverse effectiveness. Exp Brain Res. 2009;198: 113–126. doi:10.1007/s00221-009-1880-8

36. Holmes NP. The principle of inverse effectiveness in multisensory integration: Some statistical considerations. Brain Topography. Springer US; 2009. pp. 168–176. doi:10.1007/s10548-009-0097-2

37. Houben R, Dreschler WA. Optimization of the Dutch matrix test by random selection of sentences from a preselected subset. Trends Hear. 2015;19: 233121651558313. doi:10.1177/2331216515583138

38. Stevenson RA, Nelms CE, Baum SH, Zurkovsky L, Barense MD, Newhouse PA, et al. Deficits in audiovisual speech perception in normal aging emerge at the level of whole-word recognition. Neurobiol Aging. Elsevier Ltd; 2015;36: 283–91. doi:10.1016/j.neurobiolaging.2014.08.003

39. Anderson Gosselin P, Gagné J. Older adults expend more listening effort than young adults recognizing speech in noise. J Speech Lang Hear Res. 2011;54: 944–58. doi:10.1044/1092-4388(2010/10-0069)

40. Middelweerd MJ, Plomp R. The effect of speechreading on the speech-reception threshold of sentences in noise. J Acoust Soc Am. 1987;82: 2145–7. doi:10.1121/1.395659

41. Hagerman B. Sentences for testing speech intelligibility in noise. Scand Audiol. Taylor & Francis; 1982;11: 79–87. doi:10.3109/01050398209076203

42. Hochmuth S, Brand T, Zokoll MA, Castro FZ, Wardenga N, Kollmeier B. A Spanish matrix sentence test for assessing speech reception thresholds in noise. Int J Audiol. 2012;51: 536–44. doi:10.3109/14992027.2012.670731

43. Ozimek E, Warzybok A, Kutzner D. Polish sentence matrix test for speech intelligibility measurement in noise. Int J Audiol. 2010;49: 444–454. doi:10.3109/14992021003681030

44. Wagener K, Josvassen JL, Ardenkjaer R. Design, optimization and evaluation of a Danish sentence test in noise. Int J Audiol. Taylor & Francis; 2003;42: 10–7. doi:10.3109/14992020309056080

45. Lee MD, Wagenmakers E-J. Bayesian cognitive modeling: A practical course. Cambridge University Press, 2014. Cambridge University Press, New York; 2014.

46. Kruschke JK. Doing Bayesian Data Analysis. 2nd ed. Elsevier; 2015. doi:10.1016/C2012-0-00477-2

47. Plummer M. JAGS: A program for analysis of Bayesian graphical models using Gibbs sampling. Hornik K, Leisch F, Zeileis A, editors. Proceedings of the 3rd Internaitional Workshop on Disbtributed Statistical Computing. Vienna, Austria; 2003. Available: http://mcmc-jags.sourceforge.net

48. Turner BM, Forstmann BU, Wagenmakers E-J, Brown SD, Sederberg PB, Steyvers M. A Bayesian framework for simultaneously modeling neural and behavioral data. Neuroimage. Elsevier Inc.; 2013;72: 193–206. doi:10.1016/j.neuroimage.2013.01.048

49. Brooks SP, Gelman A. General Methods for Monitoring Convergence of Iterative Simulations. J Comput Graph Stat. Taylor & Francis; 1998;7: 434–455. doi:10.1080/10618600.1998.10474787

50. Gelman A, Carlin JB, Stern HS, Rubin DB. Bayesian Data Analysis, Third Edition (Texts in Statistical Science). Book. Chapman and Hall/CRC; 2013.

51. Kass RE, Carlin BP, Gelman A, Neal RM. Markov Chain Monte Carlo in Practice: A Roundtable Discussion. Am Stat. 1998;52: 93. doi:10.2307/2685466

